# Lymphocytic choriomeningitis arenavirus requires cellular COPI and AP-4 complexes for efficient replication and virion production

**DOI:** 10.1101/2023.06.27.546563

**Authors:** Owen Byford, Amelia B. Shaw, Hiu Nam Tse, Eleanor J. A. A. Todd, Beatriz Álvarez-Rodríguez, Roger Hewson, Juan Fontana, John N. Barr

**Affiliations:** School of Molecular and Cellular Biology, Faculty of Biological Sciences, University of Leeds, Leeds, LS2 9JT, United Kingdom; Astbury Centre for Structural Molecular Biology, University of Leeds, Leeds, United Kingdom; Virology and Pathogenesis Group, National Infection Service, Public Health England, Porton Down SP4 0JG, UK

**Keywords:** arenavirus, LCMV, COPI, AP-4

## Abstract

Lymphocytic choriomeningitis virus (LCMV) is a bisegmented negative-sense RNA virus classified within the *Arenaviridae* family of the *Bunyavirales* order. LCMV is associated with fatal disease in immunocompromised populations, and as the prototypical arenavirus, acts as a model for the many serious human pathogens within the *Arenaviridae* family. Here, we examined the dependence of LCMV multiplication on cellular trafficking components using a recombinant LCMV expressing enhanced green fluorescent protein in conjunction with a curated siRNA library. The screen revealed a requirement for subunits of both the coat protein 1 (COPI) coatamer and adapter protein 4 (AP-4) complexes. By rescuing a recombinant LCMV harbouring a FLAG tagged GP-1 envelope spike (rLCMV-GP1-FLAG) we showed infection resulted in marked co-localization of COPI and AP-4 component with both LCMV nucleoprotein (NP) and GP-1. Time-of-addition studies using brefeldin A (BFA), an ARF-I inhibitor that prevents formation of both COPI and AP-4 complexes, suggested these cellular components were involved in late stages of the LCMV multiplication cycle. Consistent with this finding, BFA treatment at similar late time-points resulted in a marked redistribution of NP and GP-1, and subsequent loss of COPI/AP-4 co-localization. Finally, titration of released virus within supernatant of BFA-treated cells revealed a 10-fold decrease in viral titres, greater than the 2-fold BFA-mediated reduction in NP expression. Taken together, these findings suggest COPI and AP-4 complexes are important host cell factors that are required for efficient LCMV assembly and egress.

**Importance:** Arenaviruses are rodent-borne, segmented, negative-sense RNA viruses, with several members responsible for fatal human disease, with the prototypic member LCMV being under-recognised as a pathogen capable of inflicting neurological infections with fatal outcome. Here, we assessed the impact of siRNA knockdown of host cell trafficking genes on LCMV multiplication. We reveal the requirement of host cellular COPI and AP-4 complexes for efficient LCMV multiplication, acting late in the replication cycle, at the stages of egress and assembly. Collectively, our findings improve the understanding of arenaviruses host-pathogen interactions and reveal novel cellular trafficking pathways required during infection. Moreover, this study may lead to the discovery of novel therapeutic targets for arenaviruses to prevent serious human disease.

## Introduction

The *Arenaviridae* family within the *Bunyavirales* order of segmented negative-sense RNA viruses currently compromises 56 species, which are sub-divided into four genera: *Antennavirus*, *Hartmanivirus*, *Mammarenavirus* and *Reptarenavirus*. Members of the *Mammarenavirus* genus can infect mammals [1–3] and notable species include Lassa virus (LASV) and Junín virus (JUNV), for which infections can result in haemorrhagic fever and subsequent fatality in approximately 1-2% and 30% of human cases, respectively [4, 5]. Currently, no specific antiviral therapeutics or FDA-approved vaccines exist to target any member of the *Arenaviridae* family [6]. Combined, these factors contribute to the classification of LASV, JUNV and several other arenaviruses as hazard group 4 pathogens, requiring the highest biosafety level (BSL) 4 containment for their study, which has hindered research progress.

Lymphocytic choriomeningitis virus (LCMV) is classified within the *Mammarenavirus* genus and causes a persistent infection within the common house mouse, *Mus musculus*, which acts as the primary host. Rodent-to-human transmission frequently occurs but rarely results in severe disease [7]. LCMV exhibits tropism for non-differentiated neuroblasts and in some populations, including immunocompromised patients or neonates, can lead to aseptic meningitis [8, 9]. Specifically, the LCMV Armstrong strain acts as an effective research model for more pathogenic mammarenaviruses due to shared structural and functional characteristics, and requirement for more amenable BSL-2 containment facilities for its study.

The LCMV genome comprises small (S) and large (L) segments and encodes four structural proteins using an ambi-sense transcription strategy. Following entry, the input virion-associated (vRNA) S and L segments are transcribed by the viral RNA-dependent RNA polymerase (RdRp) to generate mRNAs encoding nucleocapsid protein (NP) and RdRp, respectively. Subsequent vRNA replication yields S and L anti-genome (ag) RNAs, which act as templates for the transcription of mRNAs encoding a glycoprotein precursor (GPC) and matrix protein (Z) [10, 11]. Post-translational cleavage of GPC results in the production of N-terminal glycoprotein-1 (GP-1), C-terminal glycoprotein-2 (GP-2) and stable signal peptide (SSP), that remain associated as trimers [12–15].

Although LCMV has been studied heavily from an immunological perspective, many molecular details underpinning its multiplication are yet to be determined. It is thought that LCMV first attaches to the cell at the plasma membrane (PM) by binding to cellular receptor α-dystroglycan, although members of the Tyro3, Axl, and Mer (TAM) family and C-type lectins have also been implicated [16, 17]. LCMV is internalised, potentially via macropinocytosis due to the requirement of actin remodelling and Pak1 [18], after which virions are trafficked via multivesicular bodies (MVBs) to late endosomes. Here, following interaction with secondary receptor CD164, GP-2 mediates endosomal fusion and subsequent release of the vRNAs into the cytosol [19, 20]. The site of arenavirus RNA synthesis is unclear, with reports of newly made mRNAs located within processing bodies [21] and agRNA and vRNAs compartmentalised within specialised NP-induced organelles known as replication-transcription complexes [22]. Following translation, GPC is glycosylated within the endoplasmic reticulum (ER) then post-translationally cleaved by the resident ER signal peptidase to release the SSP, which remains associated with the GPC whilst it is trafficked to, and throughout, the Golgi [23]. Here, GPC is further cleaved by subtilisin kexin isozyme-1/site-1 protease (SKI-1/S1P) to release GP-1 and GP-2, which associate alongside SSP as trimers [24–26]. While the site of viral budding is unclear, virion assembly requires Z-mediated recruitment of endosomal sorting complex required for transport (ESCRT) components; Z is proposed to recruit RNPs via interaction with NP, and myristylation of Z is required for virion incorporation of glycoprotein trimers [27, 28].

Here, using a curated siRNA library targeting cellular proteins involved in vesicular trafficking we showed the efficient LCMV multiplication in the neuronal cell line SH-SY5Y required components of the coat protein I (COPI) and adaptor protein 4 (AP-4) complexes. Disrupting the formation of these complexes using brefeldin A (BFA) resulted in redistribution of LCMV GP-1 and NP, suggesting a role for AP-4 and COPI components in LCMV protein trafficking, with BFA time-of-addition assays indicating COPI and AP-4 complexes are required for efficient LCMV assembly and egress. Overall, this study improves our understanding of arenavirus-host interactions, which may aid in the identification of targets for effective anti-arenaviral therapies.

## Results

### Identification of the trafficking components required during LCMV multiplication

A recombinant LCMV expressing eGFP (rLCMV-eGFP) using a P2A-linked [29–32] eGFP open reading frame (ORF) appended to the NP ORF (Figure 1A) was rescued as previously described [32] for which measurement of total integrated intensity of eGFP expression (TIIE) represented a surrogate for LCMV multiplication and infection progression (Figure 1B). The role of host cell trafficking components in LCMV multiplication and egress was assessed using a focused siRNA library comprising three different siRNAs specific for each of 87 host trafficking genes. Following reverse transfection of each individual siRNA, human-origin neuronal SH-SY5Y cells were infected with rLCMV-eGFP at a multiplicity of infection (MOI) of 0.2 and live-cell images were acquired to measure TIIE at 24 hpi (Figure 1D). At this time point, rLCMV-eGFP has undergone multiple infection cycles in these cells, [32]. Thus, a significant reduction in TIIE relative to the negative control (virus plus transfection reagent only) could represent inhibition of any stage of the LCMV replication cycle encompassing binding, entry, replication, assembly or egress.Following reverse transfection of each individual siRNA, human-origin neuronal SH-SY5Y cells were infected with rLCMV-eGFP at a multiplicity of infection (MOI) of 0.2 and live-cell images were acquired to measure TIIE at 24 hpi (Figure 1C). At this time point, rLCMV-eGFP has undergone multiple infection cycles in these cells [32]. Thus, a significant reduction in TIIE relative to the negative control (virus plus transfection reagent only) could represent inhibition of any stage of the LCMV replication cycle encompassing binding, entry, replication, assembly or egress.

**Figure 1.**
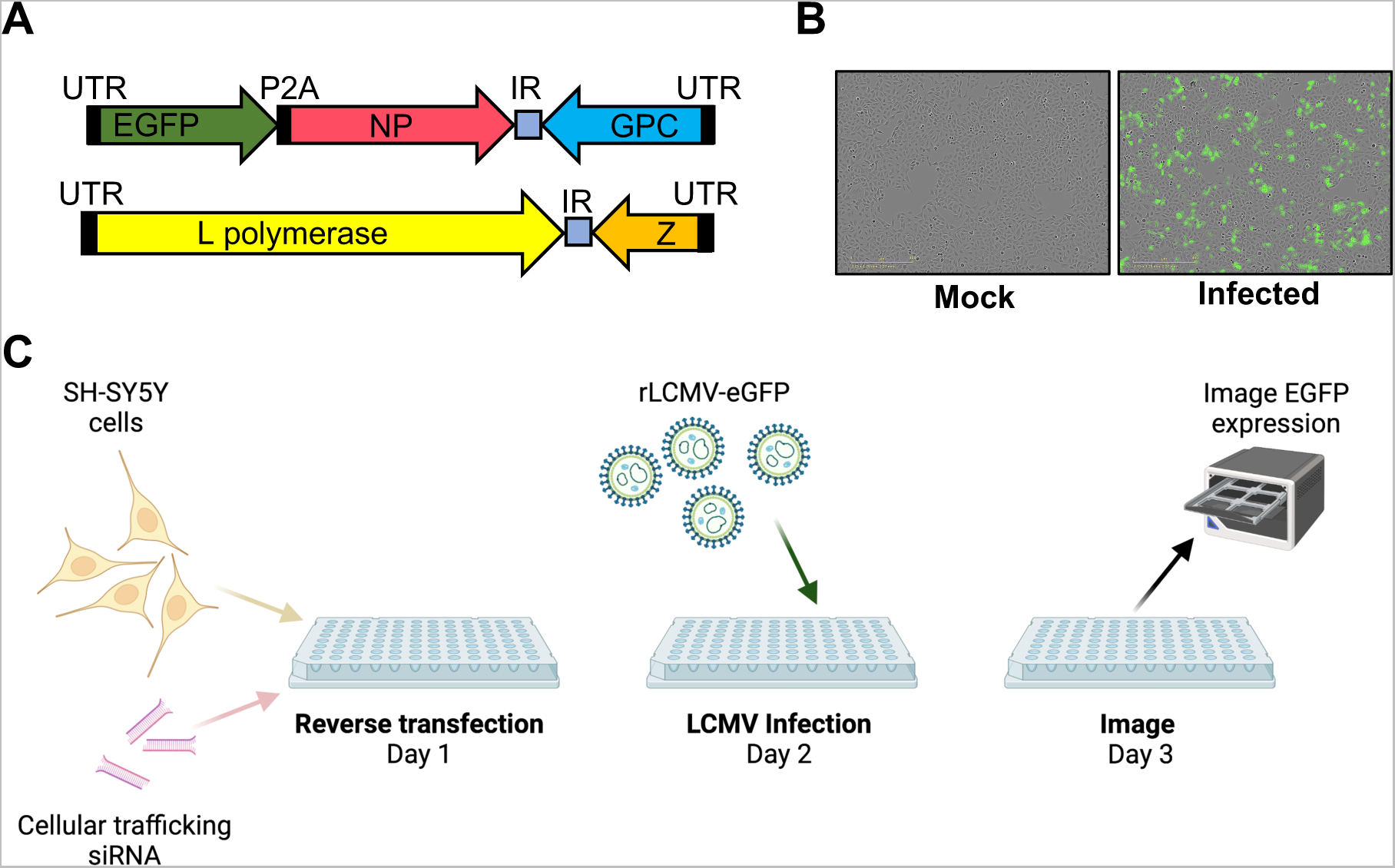
Rescue of recombinant LCMV expressing eGFP used to perform an siRNA screen to identify cellular factors involved in LCMV multiplication. (A) Schematic of rLCMV S and L segments, showing the location of the eGFP open reading frame within the S segment; UTR; untranslated region; P2A, P2A linker; IR, intergenic region. (B) Live cell images of A549 cells infected with LCMV-eGFP showing eGFP fluorescence at 24 hpi. (C) Schematic of the siRNA screen protocol.

The impact on overall multiplication and infectious virion production of each individual siRNA was tested four times (supplementary data set S1) and the 30 genes with the greatest mean TIIE reduction across all three individual siRNAs are listed in Figure 2.

**Figure 2.**
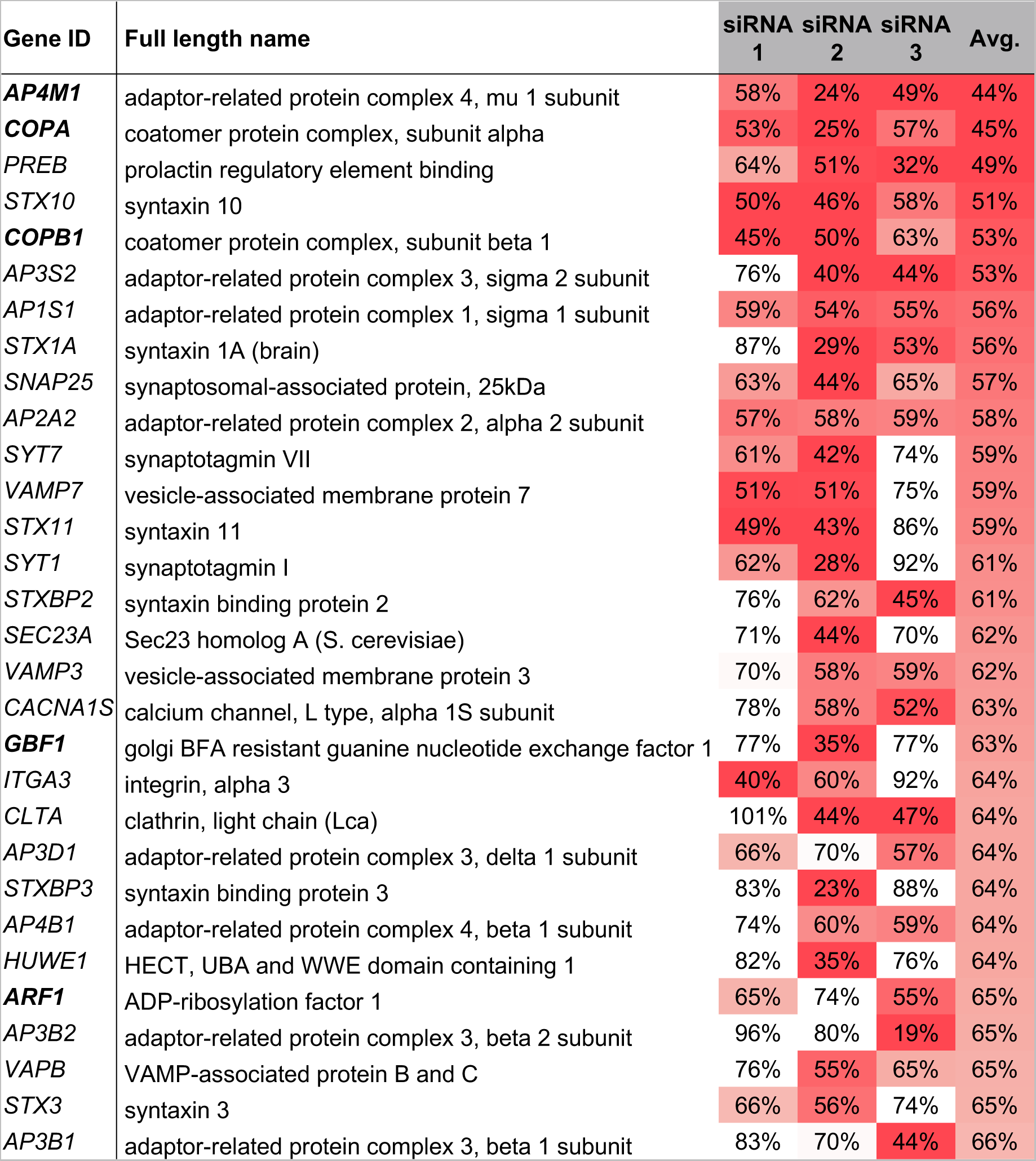
Top 30 gene targets impacting rLCMV-eGFP growth, identified using an siRNA screen of cellular factors involved in vesicle trafficking. Viral growth was measured by total integrated intensity of eGFP expression (TIIE) relative to virus plus transfection reagent control, following siRNA knockdown at 24 hpi. Percentage TIIE values are shown for three individual siRNAs against a single gene target. Each percentage value represents the mean of four experimental repeats, and the colours indicate the levels of reduction in TIIE from red (most reduction in TIIE) to white (least reduction in TIIE). Genes highlighted in boldface represent those associated with COPI and AP-4 vesicles.

### AP-4 and COPI vesicle components are required for productive LCMV infection

rLCMV-eGFP-mediated TIIE in siRNA transfected cells was most-reduced following knockdown of AP-4 component AP4M1, with a mean TIIE level at 24 hpi of 44% compared to scrambled (Figure 2; siRNA 1 = 0.5771 [P = 0.1399]; siRNA 2 = 0.2412 [P = 0.1907]; siRNA 3 = 0.4935 [P=0.0860]). In addition, knockdown of a second AP-4 component, AP4B1, resulted in a mean TIIE level of 64%. The second most-reduced TIIE level was for COPI vesicle component COPA, with 45% TIIE compared to scrambled (Figure 2; siRNA 1 = 0.5335 [P = 0.0354]; siRNA 2 = 0.2524 [P = 0.0008]; siRNA 3 = 0.5664 [P = 0.0014]). In addition, knockdown of a second COPI component, COPB1, resulted in 53% TIIE.

Interestingly siRNA knockdown of other components required indirectly to form both AP-4 and COPI vesicles, namely the small GTP-binding protein ADP-ribosylation factor 1 (ARF-1) and Golgi BFA-resistant guanine nucleotide exchange factor 1 (GBF-1) (30), also resulted in significantly reduced TIIE levels. GBF1 is a guanine exchange factor (GEF) responsible for activation of ARF-1, which then binds GTP to allow recruitment of intact COPI heptameric complexes and their subsequent anchoring within membranes [33]. The recruitment of AP-4 vesicles to the TGN is also regulated by ARF-1, a process mediated by direct interaction between AP4M1 and ARF-1, following its activation [34]. Taken together, these findings suggested important roles for both AP-4 and COPI complexes in the LCMV multiplication cycle.

### Generation of a recombinant infectious LCMV bearing a FLAG tagged GP-1

AP-4 and COPI complexes are known to localise to the Golgi [34–37] given that LCMV GPC is known to be proteolytically-cleaved and post-translationally modified within this compartment [26], we wished to develop a tool to simultaneously assess GP distribution alongside NP during virus infection.

To achieve this, we generated rLCMV with a FLAG epitope tag inserted within GP-1 (rLCMV-GP1-FLAG). The site for FLAG insertion was rationally selected by identification of regions of high sequence variation between various OW arenavirus members (Figure 3A) and a SWISS-MODEL prediction of the GP-1 structure, which revealed the selected site was within a flexible and surface exposed region, thus a suitable candidate for detection by immunofluorescence (IF). Subsequent virus rescue was confirmed by western blotting (Figure 3B) and using an immunofluorescent focus forming assay utilising NP and FLAG (GP-1) antibody staining, which revealed the resulting virus rLCMV-GP1-FLAG was infectious, the FLAG epitope was accessible to antibody detection, and yielded viral titres of around 10^5^ pfu/mL (Figure 3C)

**Figure 3.**
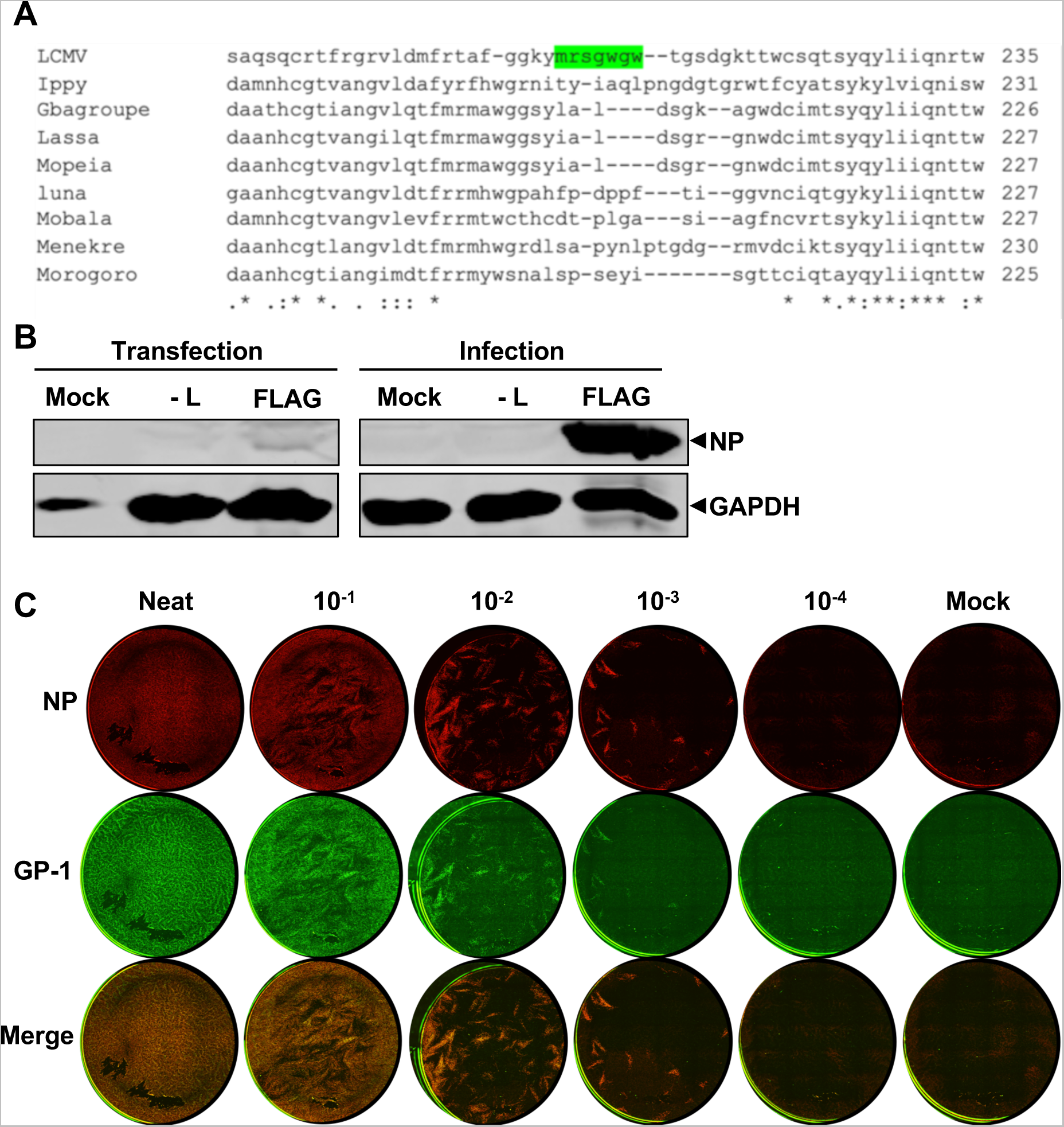
Generation of a recombinant LCMV expressing a FLAG-tagged GP-1. (A) Partial GP-1 sequence alignment of selected Old-World arenaviruses identifying a region of low conservation as a potential FLAG tag insertion site (green). (B) Western blot analysis of transfected BSR-T7 and infected BHK-21 cell cultures confirming LCMV-GP1-FLAG rescue, using antisera specific for LCMV NP and GAPDH as loading control. (C) Immunofluorescence focus forming assay for rLCMV-GP1-FLAG showing foci by staining at 3 days post infection for LCMV NP, GP-1 (FLAG) alongside merged channels.

### Analysis of LCMV NP and GP-1 localization

IF analysis of rLCMV-GP1-FLAG infected A549 cells at 24 hpi using NP and FLAG antisera revealed NP and GP-1 distribution was distinct; while NP was distributed widely in perinuclear regions with occasional dense puncta, GP-1 was mostly localised on one side of the nucleus, reminiscent of Golgi distribution (Figure 4A). To confirm Golgi localisation of GP-1, we performed IF using antisera specific for both FLAG and the *cis*-Golgi marker GM130, for which many of the corresponding peak intensities corresponded, quantified by line scan analysis across a distance of approximately 20 µm (Figure 4B). This confirmed previous localization studies that used transient LCMV GPC expression [38], although here, we report for the first time the cellular location of GP-1 expressed from an infectious LCMV.

**Figure 4.**
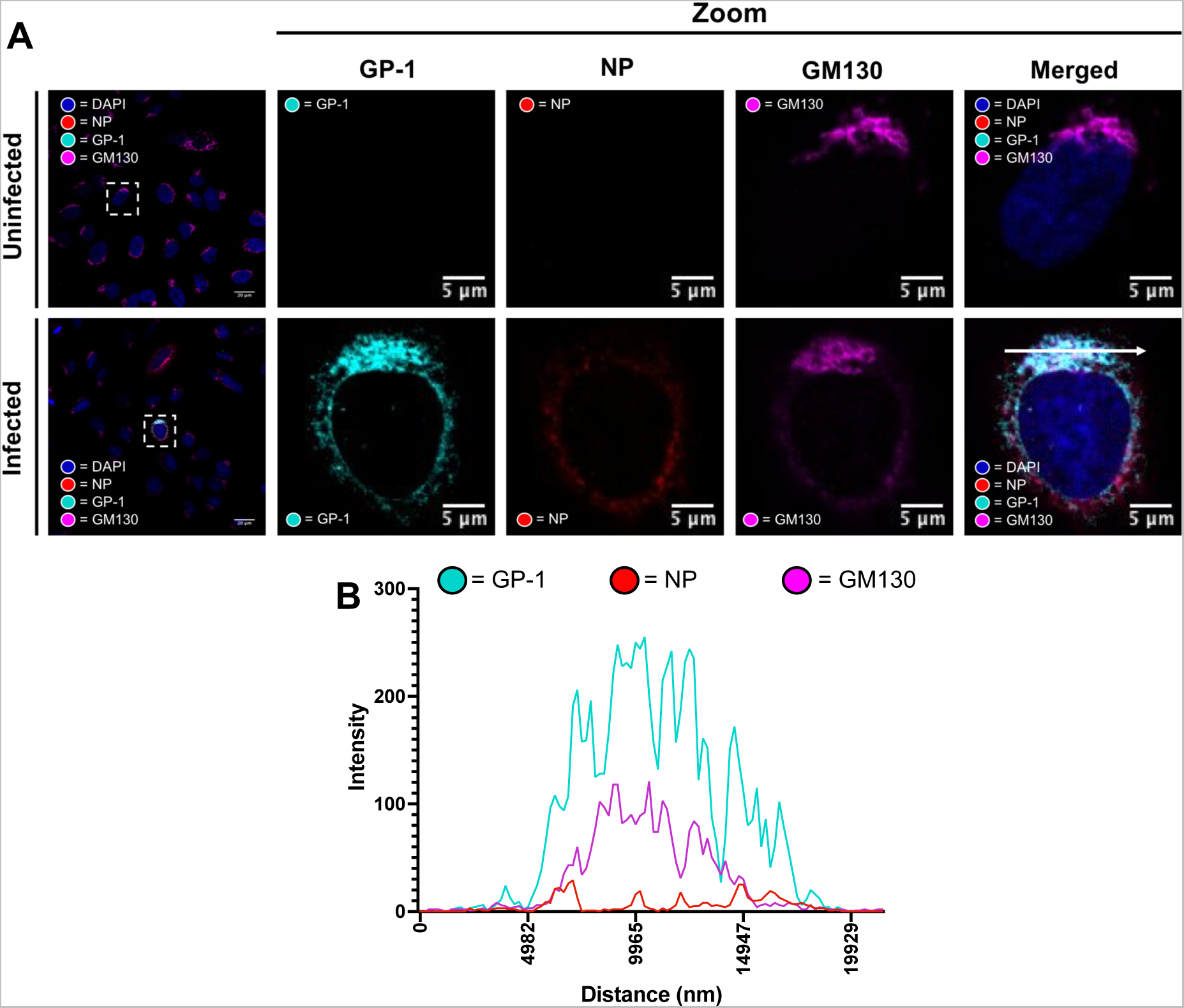
GP-1 expressed from rLCMV-GP1-FLAG co-localises with Golgi marker GM130. (A) Confocal microscopy of uninfected and rLCMV-GP1-FLAG infected cells at an MOI of 0.1 at 24 hpi. A549 cells were stained with antisera specific for GM130 (magenta), GP-1 (cyan), and NP (red) alongside DAPI. A zoomed image for both uninfected and infected (white boxes) for all channels is shown, alongside merged channels. (B) Line scan for the region highlighted in the zoomed merged image in panel A (scan line represented by a white arrow), showing intensities of channels corresponding to NP, GP-1 and GM130 across a ∼ 20 µm distance.

### rLCMV infection results in redistribution and upregulated expression of COPI complex components

Using rLCMV-GP1-FLAG, we next examined the spatial proximity of NP and GP-1 alongside components of COPI complexes.

As expected, COPA staining within uninfected cells presented as discrete puncta with characteristic ER/Golgi distribution (Figure 5A-B). In contrast, within rLCMV-GP1-FLAG infected cells, while some COPA staining also exhibited this discrete ER/Golgi localisation, its distribution was expanded outside perinuclear regions, extending throughout the cytoplasm. This redistributed COPA staining closely corresponded with that of NP, which is strikingly represented in the merged image and associated line scan (Figure 5C), suggesting NP and COPA may directly interact to cause this redistribution.

**Figure 5.**
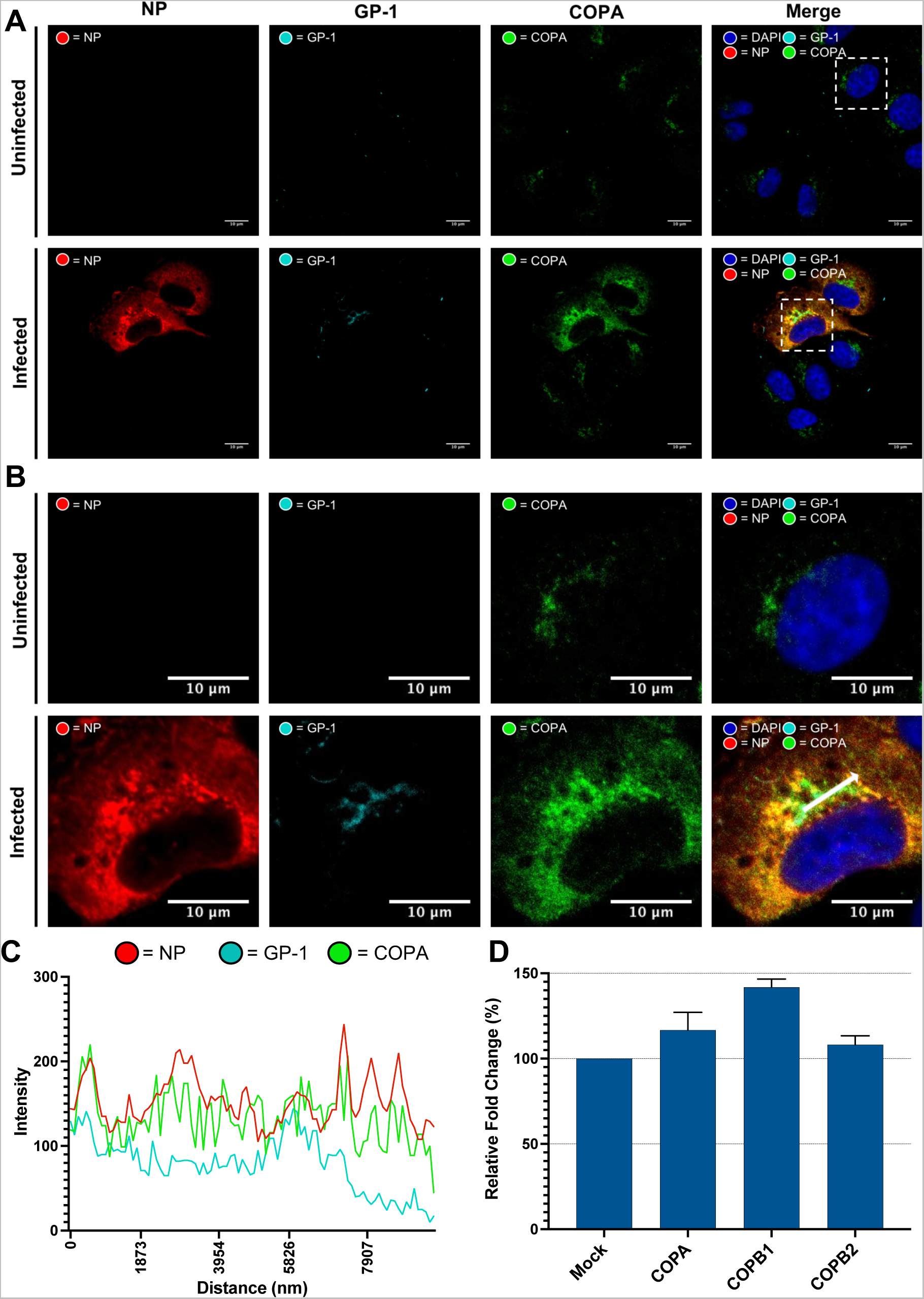
LCMV NP and the COPA component of COPI complexes co-localize in LCMV infected cells. (A) Confocal microscopy of uninfected and rLCMV-GP1-FLAG infected cells 15 hpi, MOI of 0.1. A549 cells were stained with antisera specific for COPA (green), GP-1 (cyan), and NP (red) alongside DAPI. (B) A zoomed image for both uninfected and infected (white boxes) for all channels is shown, alongside merged channels. (C) Line scan for the region highlighted in the zoomed image in panel B (scan line represented by a white arrow), showing intensities of channels corresponding to NP, GP-1 and COPA across a ∼10 µm distance. (D) Quantitation of COPA, COPB1 and COPB2 mRNA expression in uninfected and LCMV infected (MOI 1) cells via qRT-PCR. Four experimental repeats were performed for each individual gene, with the variance from the mean shown with error bars.

Co-staining with FLAG anti-sera revealed GP-1 occupied a restricted area within that occupied by COPA, with many of the peak intensities of GP-1 and COPA being closely aligned, as shown by line scan analysis (Figure 5C). This GP-1 and COPA colocalization was consistently imaged across all infected cells analysed (Supplementary Figure S1A). As with our previous observations, described above (Figure 4A and B), NP and GP-1 occupied distinct regions, and the positions of their peak intensities or general abundance did not precisely correspond.

Alongside altered COPA localisation, rLCMV-GP1-FLAG infection also appeared to result in increased COPA abundance, as judged by higher intensity of IF staining (Figure 5A and B). To quantify this apparent increased expression, qRT-PCR was used to compare COPA and COPB1 mRNA levels in LCMV-infected versus mock cells (Figure 5D), which showed a 16% and 41% increase, respectively, corroborating the observed increased expression in the corresponding IF images.

Taken together, these findings suggest a close association between COPA and NP; and also, between COPA and GP-1, consistent with a role for COPI in LCMV NP and GPC processing, as well as significant upregulation of COPI components following LCMV infection.

### LCMV utilises AP-4 complexes and upregulates AP4M1 expression

As with components of the COPI complex described above, rLCMV-GP1-FLAG allowed examination of the spatial proximity of NP and GP-1 alongside the AP4 complex. Staining uninfected cells with antisera for AP4 subunit AP4E1 revealed staining (Figure 6A-B) consistent with its previously characterised distribution within the *trans*-Golgi network [36, 37]. In contrast, within rLCMV-GP1-FLAG infected cells, while some AP4E1 localised to punctate regions, the majority was located diffusely throughout the cytoplasm, suggestive of re-distribution.

**Figure 6.**
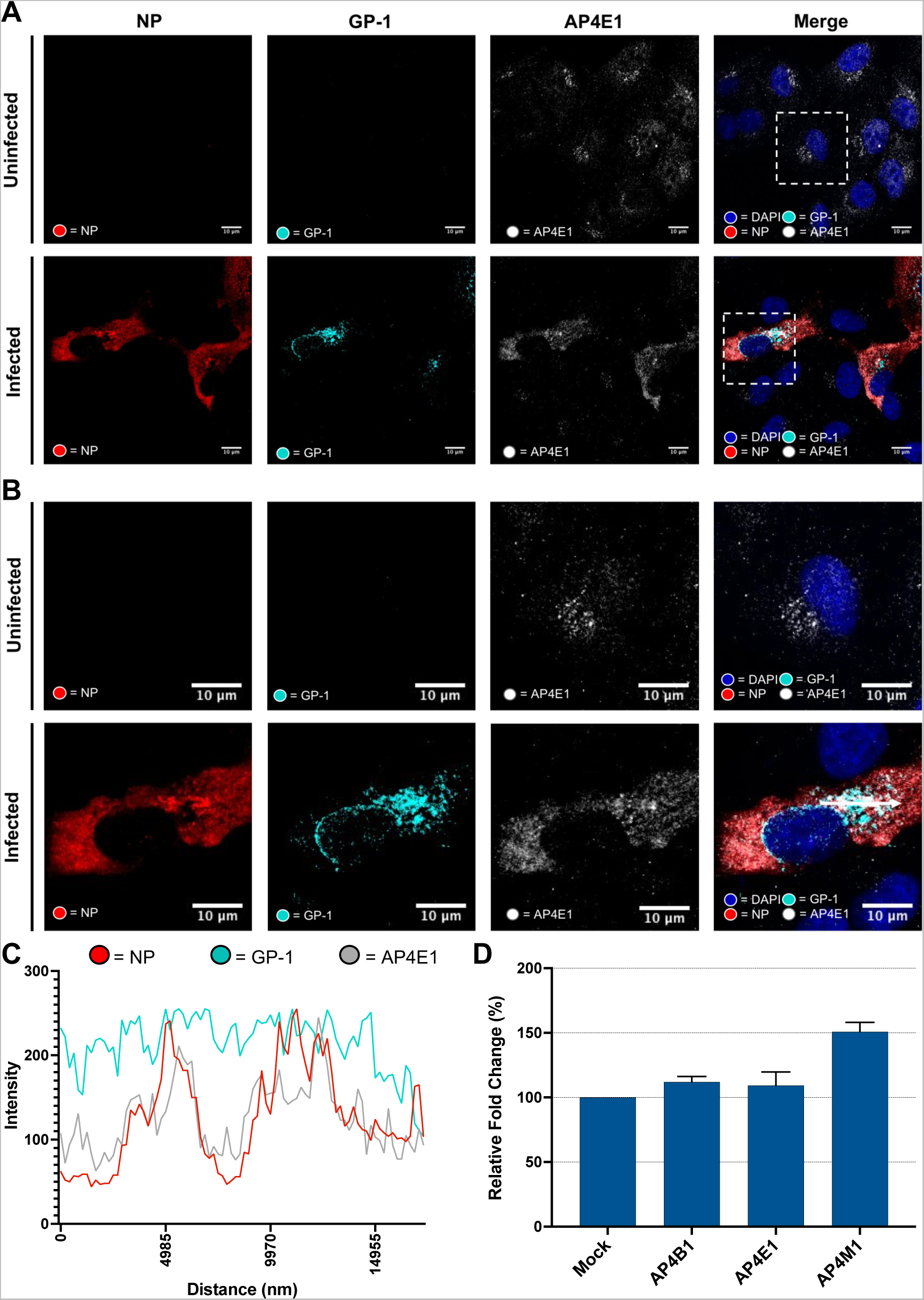
LCMV NP and the AP4E1 component of AP-4 complexes co-localize in LCMV infected cells. (A) Confocal microscopy of uninfected and rLCMV-GP1-FLAG infected cells at 15 hpi, MOI of 0.1. A549 cells were stained with AP4E1 (white), GP-1 (cyan), and NP (red) and DAPI. (B) A zoomed image for regions of both uninfected and infected cells (white boxes) for all channels is shown, alongside merged channels. (C) Line scan analysis for the region highlighted in the merged zoomed image shown in panel B (scan line represented by a white arrow) showing intensities of channels corresponding to NP, GP-1 and AP4E1 across a ∼17 µm distance. (D) Quantitation of mRNA expression levels for AP4B1, AP4E1 and AP4M1 in uninfected and LCMV infected (MOI 1) cells via qRT-PCR. Four experimental repeats were performed for each individual gene, with the variance from the mean shown with error bars.

Staining rLCMV-GP1-FLAG infected cells with NP antisera revealed close co-localisation with AP4E1, suggesting NP was responsible for the observed AP4E1 redistribution (Figure 6C). In contrast, while GP-1 staining occupied a portion of that occupied by AP4E1/NP, its distribution was less extensive and more reminiscent of Golgi, as described above. To gain a better overall representation of GP-1, NP and AP4E distribution, colocalization of all these components was imaged in multiple different rLCMV-GP1-FLAG infected cells (Supplementary Figure S1B), which revealed a consistent outcome.

Alongside altered AP4E1 localisation, rLCMV-GP1-FLAG infection appeared to result in increased AP4E1 abundance, as judged by higher intensity of AP4E1 staining. To quantify this apparent increase, qRT-PCR analysis was used (Figure 6D) to quantify AP4B1, AP4E1 and AP4M1 mRNA expression from uninfected and infected cells (MOI 1). While expression levels of AP4B1 and AP4E1 were slightly increased, AP4M1 mRNA expression was found to be upregulated by 50%.

### Abrogation of AP-4 and COPI complex formation using BFA disrupts LCMV gene expression

BFA inhibits activation of ARF-1, blocking formation of COPI and AP-4 complexes [39, 40]. We first confirmed BFA was non-cytotoxic for human-origin A549 cells in concentrations up to 5,000 ng/mL (Supplementary Figure 2A). We then confirmed the expected cellular changes for A549 cells by treatment with BFA followed by staining with antisera specific for COPA and AP4E1 sub-components (Supplementary Figure 2B). Under normal cellular conditions, both COPA and AP4E1 localised in discrete perinuclear regions, consistent with Golgi/TGN localisation. However, upon addition of 5000 ng/mL BFA for 1 hour both COPA and AP4E1 staining became diffusely distributed throughout the cytoplasm.

Next, to confirm the involvement of ARF-1 and GBF-1 in LCMV multiplication, we examined the effect of BFA on LCMV gene expression. A549 cells were pre-treated for 45 min with BFA at various concentrations prior to infection with rLCMV-WT at an MOI of 0.1. At 24 hpi, cell lysates from rLCMV-WT infections were harvested and virus multiplication analysed by western blotting utilising NP antisera (Figure 7A). Influenza A virus (IAV) was utilised as a positive control, with BFA treatment known to result in diminished, but not abrogated, IAV gene expression and infectious virion assembly [41, 42]. Infections were performed in triplicate, with respective NP abundances analysed via densitometry (Figure 7B). For LCMV, this analysis showed a marked reduction in LCMV NP abundance at BFA concentrations between 50 and 100 ng/mL, after which NP expression was diminished but remained constant at approximately 40% that of untreated controls, alongside no detectable change in GAPDH expression, and no cellular toxicity (Supplementary Figure 2A). IAV NP expression showed a similar decrease in abundance across the same BFA concentration range, as expected [41, 42].

**Figure 7.**
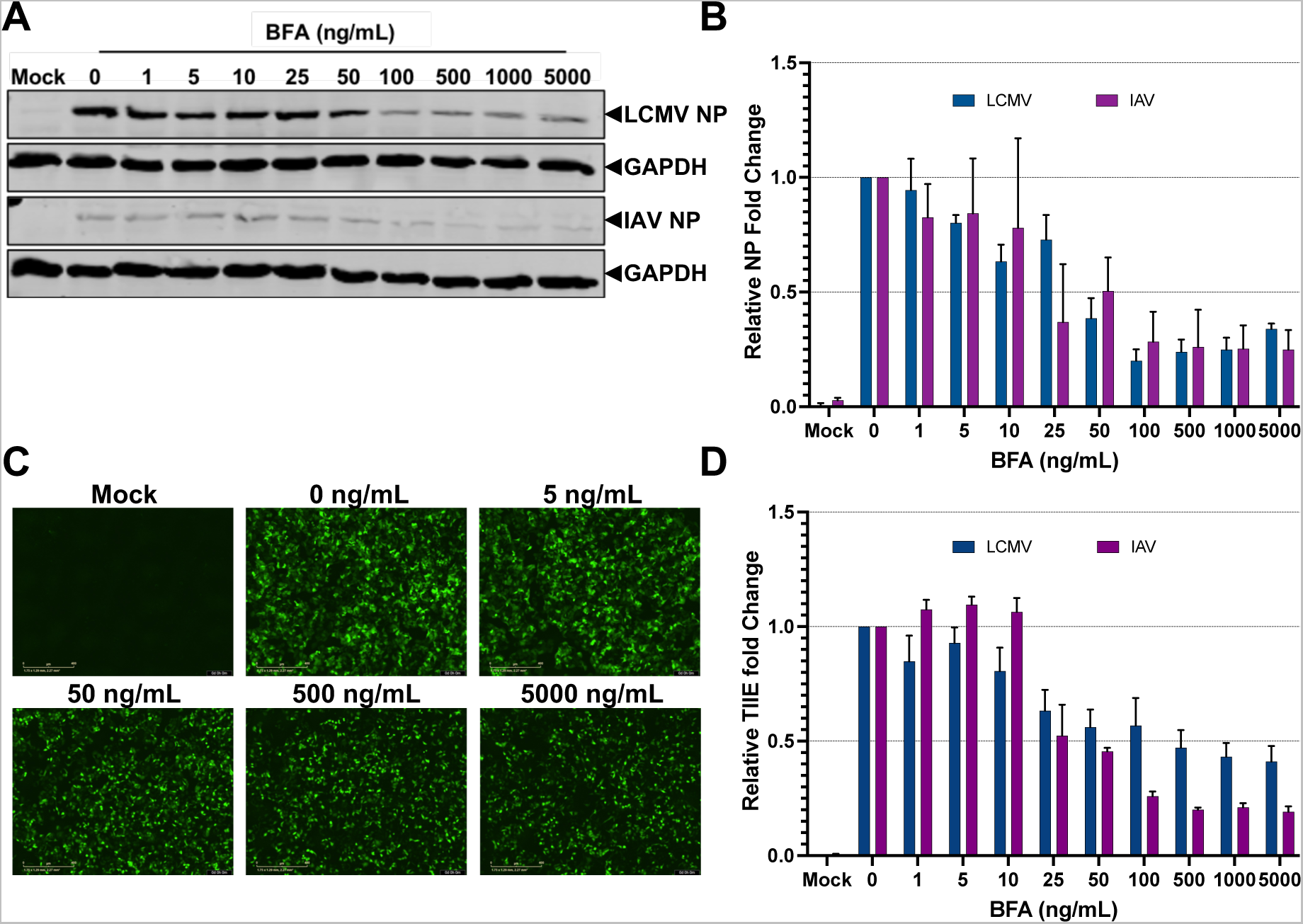
LCMV multiplication is inhibited following BFA treatment suggesting a requirement for ARF-1 and GBF-1. (A) Representative western blot analysis of A549 cells pre-treated with BFA 45 minutes prior to WT LCMV and IAV infection at an MOI of 0.1, with lysates harvested at 24 hpi. Samples were probed using antisera for their respective NPs (LCMV NP, IAV NP) and GAPDH as a loading control. (B) Densitometry histograms of the relative fold change in NP expression for LCMV (blue) and IAV (green) of cultures shown in panel A. Expression was quantified for three independent experimental repeats, with error bars representing deviation from the mean. (C) Representative fluorescent microscopy images of A549 cells pre-treated with BFA at indicated concentrations (0, 5, 50, 500 and 5000 ng/mL) and infected with rLCMV-eGFP at an MOI of 0.1 or uninfected (mock). (D) Histograms of the relative fold change in normalized TIIE in A549 cells pre-treated with BFA, and then infected with rLCMV-eGFP (blue) and rIAV-eGFP (green) at an MOI of 0.1. TIIE expression was quantified for three independent experimental repeats, with error bars representing deviation from the mean.

To corroborate these findings, BFA pre-treated A549 cultures were also infected with rLCMV-eGFP to assess the impact of BFA on eGFP expression, as a marker for viral gene expression. As expected, increasing concentration of BFA caused a decrease in TIIE expression with results closely corresponding to those for NP expression (Figure 7C-D), at approximately 50% that of untreated controls at higher BFA concentrations. As above, as a positive control for BFA efficacy, the TIIE measured from an eGFP-expressing variant of IAV showed an expected reduction at higher BFA concentrations.

### Time-of addition experiments show BFA does not affect early stages of the LCMV replication cycle

The finding that LCMV multiplication was diminished by BFA pre-treatment and siRNA-mediated COPA and AP4M1 knock down suggests COPI and AP-4 complexes play important roles in LCMV multiplication. To determine the stage of the LCMV replication cycle at which these complexes are required, we performed time-of-addition experiments. Cells were treated with BFA at time points of 0, 3, 6 and 9 hpi post infection, with times chosen to target temporally-distinct stages of the LCMV multiplication cycle; internalisation is complete at 3 hpi, 6 hpi signifies the onset of gene expression, with 9 hpi corresponding to when assembled virions are first released [32]. All infected cultures were lysed at 24 hpi, with NP expression examined by western blotting (Figure 8A), quantified by densitometry (Figure 8B).

**Figure 8.**
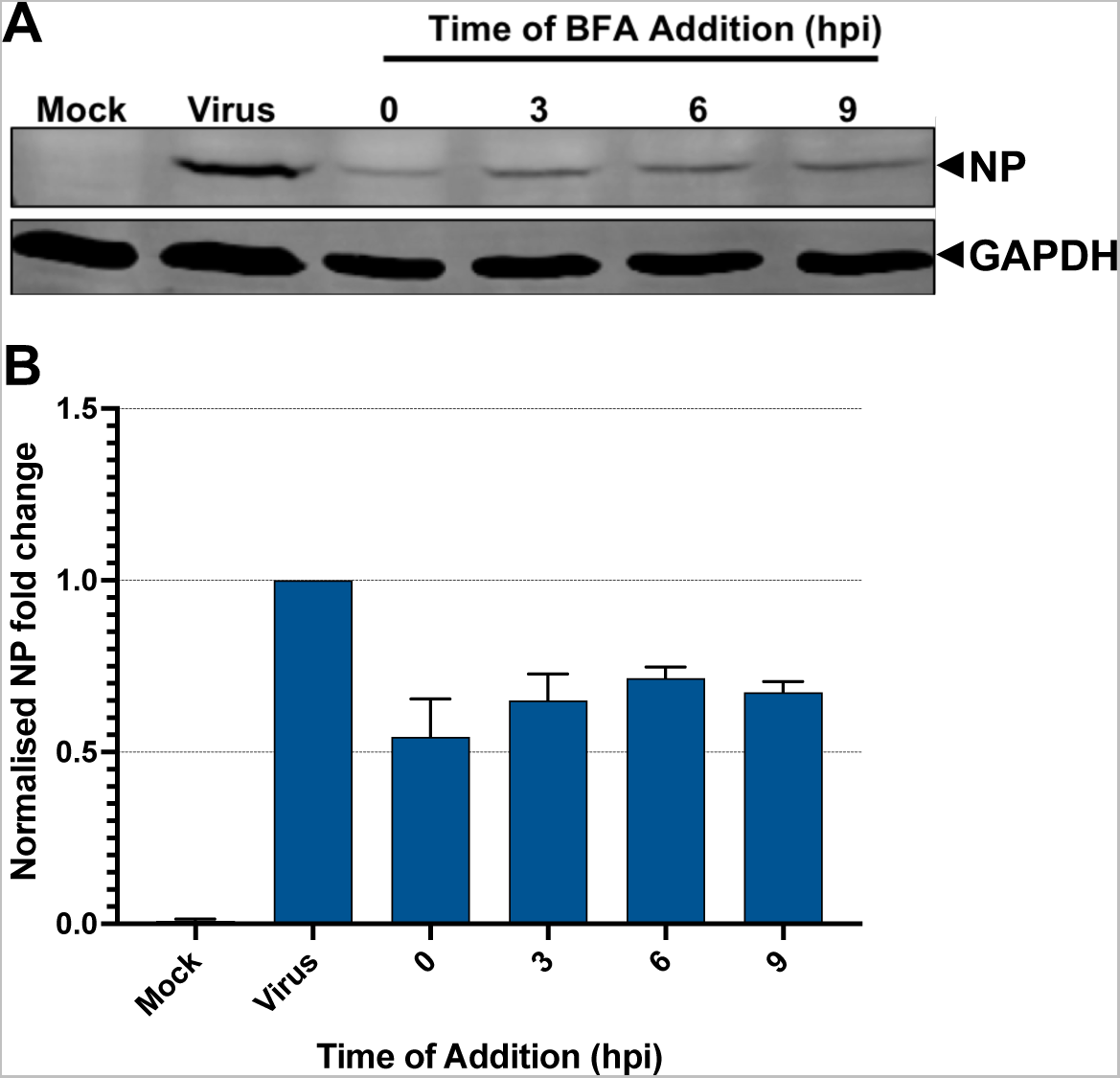
Time of addition assay indicates disruption of AP-4 and COPI complex formation using BFA does not affect early stages of the LCMV multiplication cycle. A549 cells were infected with rLCMV-WT at an MOI of 0.1 and a time of addition assay was performed, with BFA added at the indicated hpi and remaining present for the duration of infection. At 24 hpi, cell lysates were collected and subject to western blot analysis. (A) Representative western blot analysis of the time of addition assay, with samples probed using antisera for LCMV NP and GAPDH as a loading control. (B) Densitometry histogram of the relative fold-change of NP expression quantified for three independent experimental repeats with error bars showing deviation from the mean.

In line with previous results (Figure 7A and B), BFA treatment of WT-LCMV infected cells at 0 hpi, thus present for the entire replication cycle, resulted in NP expression levels of approximately 50% compared to untreated cultures (Figure 8A and B). Addition of BFA at 3 hpi, when internalisation is complete, resulted in a statistically indistinguishable reduction in NP expression levels compared to 0 hpi treated cultures. The finding that NP expression was not significantly changed when BFA was added either before (0 hpi) or after internalisation and entry stages (i.e. 3, 6 or 9 hpi time points) showed BFA did not influence the early stages of the LCMV replication cycle. Furthermore, the finding that BFA addition at 6 or 9 hpi, time points that are post-entry and encompass the onset of gene expression, resulted in diminished NP expression levels statistically indistinguishable to that of 0 hpi treated cultures. Taken together, these findings suggest BFA influences post-entry stages of the LCMV multiplication cycle, either at the stages of gene expression and/or assembly and egress.

### BFA disproportionately affects the assembly and egress of LCMV

As BFA treatment showed no statistically significant effect when added at early stages of the LCMV replication cycle, this implicated a role for BFA during later stages, such as gene expression or virion assembly and egress. To test this, A549 cells were pre-treated with BFA then infected with rLCMV-eGFP at an MOI of 0.1. At 24 hpi, supernatants were collected viral titres quantified by fluorescence focus forming assay, which showed that increasing concentrations of BFA reduced the production of infectious LCMV virions in a step-wise manner (Figure 9A) with a BFA concentration of 5000 ng/mL resulting in a 10-fold reduction in virus titre compared to untreated (Figure 9B).

**Figure 9.**
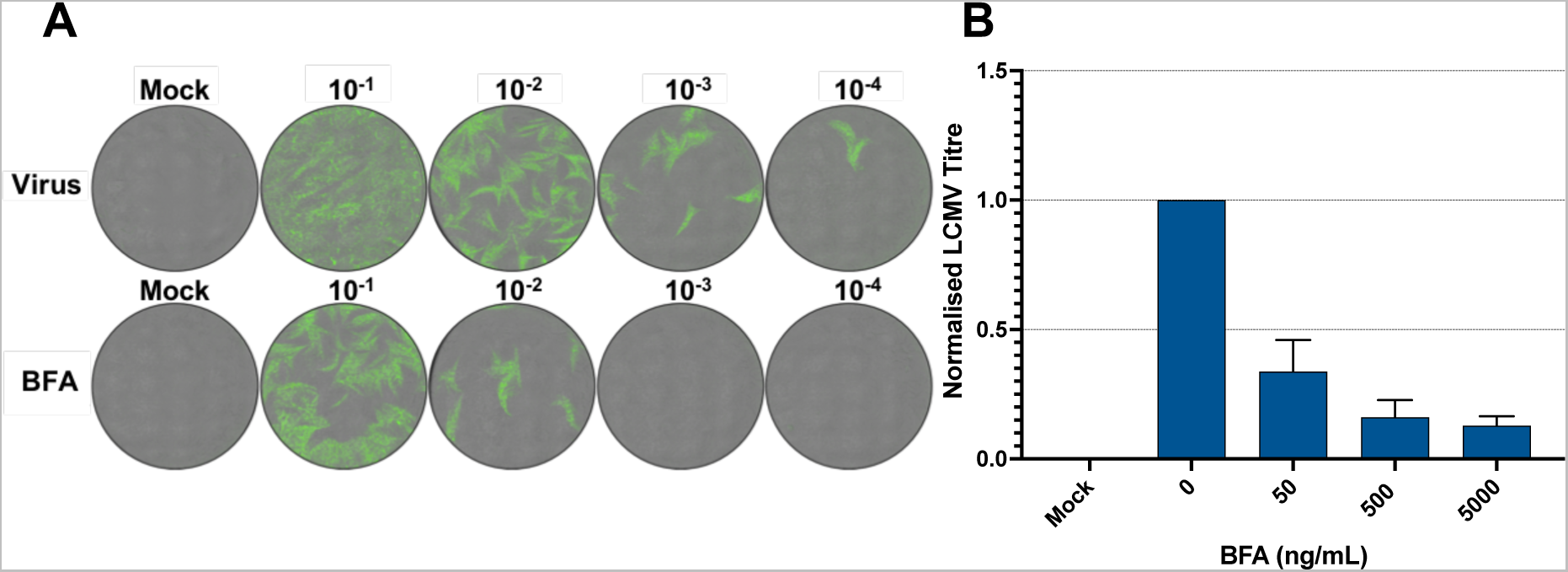
BFA treatment of rLCMV-eGFP infected cells during later stages of infection impacts infectious virus release. (A) A549 cells were pre-treated with BFA at the given concentrations for 45 mins, and then infected with rLCMV-eGFP at an MOI of 0.1. At 24 hpi, viral supernatants were harvested, serially diluted, and titres calculated via focus forming assay. Representative whole well focus-forming assay images are shown, with dilutions of 10^-1^, 10^-2^, 10^-3^ and 10^-4^. (B) Histogram representing the relative fold change in normalized LCMV titre for mock and infected cells pre-treated with BFA at indicated concentrations (0, 50, 500 and 5000 ng/mL). Titration was calculated for three independent experimental repeats, with error bars representing deviation from the mean.

In view of our previous findings that BFA treatment at 5000 ng/mL resulted in an approximately 50% reduction in NP expression (Figure 7A), a 10-fold drop in infectious virion production suggested a dual role for BFA treatment, a negative influence on LCMV gene expression but also a disproportionate effect on the assembly or release of infectious LCMV.

## Discussion

Here, we used rLCMV-eGFP for identification of host cell trafficking factors required for LCMV multiplication, with measurement of virus-specific eGFP expression allowing high-throughput screening of siRNA-mediated gene knock down of host cellular genes on various stages of the LCMV multiplication cycle.

The siRNA screen comprised 261 siRNAs, specific for 87 target genes, and revealed an important role for multiple cellular factors, of which multiple components of AP-4 and COPI coatomer complexes were highly ranked, judged by the impact of expression knock down on virus-specific eGFP expression (Figure 1). Corroborating the importance of COPI and AP-4 components, ARF-1, and GBF-1, required by both AP-4 and COPI complexes for anchorage to Golgi membranes [43], were also high-ranking in terms of LCMV gene expression knock down. AP4M1 is one of four components of the AP4 complex that perform native cellular roles in trafficking of cargo proteins from the TGN to the plasma membrane [37], whereas COPA is one of seven components of the COPI complex, with well-characterised involvement in Golgi-ER retrograde transport and intra-Golgi trafficking, but also expression of cell surface receptors and maturation of early endosome to autophagic vacuoles [44]. To gain a better understanding of AP-4 and COPI involvement in the LCMV multiplication cycle, we generated a recombinant LCMV mutant rLCMV-GP1-FLAG containing a FLAG-tagged epitope within GP-1, which alongside our existing NP antisera allowed the spatial cellular distribution of these viral components to be examined alongside AP-4 and COPI complexes within virus-infected cells. First, using this virus we showed LCMV GP-1 was predominantly located within the Golgi compartment. This was distinct from NP staining which was dispersed more widely throughout the cytoplasm, while also sharing some Golgi localization.AP4M1 is one of four components of the AP4 complex that perform native cellular roles in trafficking of cargo proteins from the TGN to the plasma membrane [37], whereas COPA is one of seven components of the COPI complex, with well-characterised involvement in Golgi-ER retrograde transport and intra-Golgi trafficking, but also expression of cell surface receptors and maturation of early endosome to autophagic vacuoles [44]. To gain a better understanding of AP-4 and COPI involvement in the LCMV multiplication cycle, we generated a recombinant LCMV mutant rLCMV-GP1-FLAG containing a FLAG-tagged epitope within GP-1, which alongside our existing NP antisera allowed the spatial cellular distribution of these viral components to be examined alongside AP-4 and COPI complexes within virus-infected cells. First, using this virus we showed LCMV GP-1 was predominantly located within the Golgi compartment. This was distinct from NP staining, which was dispersed more widely throughout the cytoplasm, while also sharing some Golgi localization.

Next, we examined the cellular localisation of AP4E1, NP and GP-1, which showed virus infection led to redistribution of AP4E1 away from its characteristic *trans*-Golgi location, to occupy regions of the cell where GP-1 and in particular NP was most abundant, suggestive of a close interaction between these components. Selection of AP-4 cargo is known to be mediated through binding of a YXXØ or dileucine based motifs [(D/E)XXXL(L/I) and (LL/LI)] within the cytoplasmic tail of cargo proteins to the AP4M1 subunit [37, 45]. LCMV NP has three YXXØ and eleven LCMV dileucine based motifs; GPC has six YXXØ and ten dileucine based motifs, although none lie within the predicted cytoplasmic tails, despite these tail domains being crucial in GP-1/GP-2 trafficking to the plasma membrane [46]. One possibility is that NP-mediated redistribution of AP-4 could be mediated by components of the GPC, for instance during transport of partially assembled RNPs in association with GPC components to a plasma membrane assembly site.

Currently there are many reports of viral subversion of members of the adapter protein complexes (AP1-5), although with few reports of viral roles specifically for the AP-4 complex [38, 47]. In the case of Hepatitis C virus (HCV), mutational studies showed that the HCV NS2 protein binds to the AP4M1 subunit via its dileucine motifs. When these motifs were mutated, HCV RNA replication was not impaired, yet extracellular viral titres were reduced [45]. This study also implicated AP-4 in promoting direct cell-to-cell spread [45]. Another example of viral AP-4 subversion is for Epstein-Barr virus (EBV) of the *Herpesviridae* family, which encodes a transmembrane glycoprotein BMRF-2 that has previously been shown to interact with the µ subunit of AP-4, permitting BMRF-2 trafficking from the TGN to the basolateral plasma membrane and thus EBV egress [48].

While further work is required in order unpick the role of AP-4 within the LCMV replication cycle, here, we report the upregulation of AP4M1 during LCMV infection (Figure 5D). In the absence of virus infection, the upregulation or disruption of AP-4 components can lead to a range of neurological disorders including aberrant ocular development, Alzheimer’s disease, cerebral palsy and congenital spastic tetraplegia [49–54]. This is thought-provoking considering that LCMV symptoms are mainly neurological, especially in foetal cases where the virus infects the brain and retina, leading to substantial injury and permanent dysfunction [9].

This study also focussed on the COPI coatomer complex, with knock down of two of its substituent components, COPA and COPB1, resulting in an approximately 50% reduction in LCMV-mediated eGFP expression. Although components of COPI complexes have previously been described as important for the LCMV replication [55], this is the first report of COPI component involvement in a neuronal cell culture system that is relevant to the LCMV replication cycle. As described above for AP-4, using rLCMV-GP1-FLAG allowed examination of GP-1 and NP localization in relation to COPI components. This analysis revealed a high degree of colocation between COPA and NP in both dense puncta and also a more dispersed cytoplasmic distribution, as well as close co-localization between GP-1 and COPA, although within a more restricted region, characteristic of the Golgi. Taken together these findings are entirely consistent with COPI components playing important roles in the virus replication cycle, corroborating the results of the siRNA knock down analysis.

Also supportive of a role for COPI in LCMV multiplication was the finding that BFA pre-treatment of infected cells resulted in 50% reduced virus gene expression, with BFA time-of-addition studies suggesting reduced NP yield is due to COPI involvement in a post-entry step. Together, these findings are consistent with COPI components being involved in establishment or efficiency of virus replication centres, where viral gene expression occurs, and for bunyaviruses such factories are localised around a modified Golgi compartment [56], also consistent with COPA and NP co-localization, and to some extent also GP-1. The finding that BFA addition results in a 10-fold reduction in infectious virion production, disproportionate to the 2-fold reduction in gene expression, may also indicate a role for COPI complexes in the other late stages of the virus replication cycle, such as protein processing and modification, virion assembly or egress. It is well-established that LCMV GPC is cleaved to GP-1/GP-2/SSP trimers by host cell protease SKI-1/S1P within the Golgi [26] and so one possibility is that canonical COPI complexes are utilised for intra-Golgi and retrograde Golgi-ER trafficking [57] that may be required for efficient GPC proteolytic processing. The use of siRNA mediated depletion or BFA treatment has provided many examples where COPI components are required for efficient virus multiplication [44], with many examples from within the *Bunyavirales* order. We have previously reported a reliance on COPI complexes and ARF-1 involvement for the bunyavirus Hazara virus, which is a species within the *Nairoviridae* family [42], a group that share many genetic and functional characteristics with the arenaviruses [58]. We showed Hazara virus requires COPI complexes in both ARF-I dependent and ARF-I independent processes, at early and late stages of the multiplication cycle [42], respectively. Particularly prominent was an influence of BFA on infectious virus production, similar to our current findings for LCMV. In addition to this, and likely related to COPI complex involvement, Uukuneimi virus, a member of the *Phenuiviridae* family within the *Bunyavirales* order, has been shown to require GBF-1 activation, and thus ARF-1 activation, for viral replication and particle assembly [59]. It is therefore possible that COPI complex involvement is a common requirement for all bunyaviruses. Whether this relates to a critical reliance on an intact Golgi compartment for formation of the bunyaviral factory, or a more general requirement for the formation of other lipid-based compartments that depend on COPI complex activity is an interesting topic for future research [60, 61].

## Materials and methods

### Plasmid design

Plasmids for rLCMV-WT and rLCMV-eGFP viral rescue were previously generated and described [32]. rLCMV-GP1-FLAG plasmid was generated by first performing sequence alignment of GP-1 alongside closely related OW arenaviruses to select a location for FLAG tag insertion. The predicated structure of GP-1 for rLCMV-GP1-FLAG was investigated using SWISS-MODEL and mapped alongside the solved LCMV GP-1 structure (REF), which showed the FLAG insertion site to be within an unsolved flexible loop of GP-1, likely accessible for antibody recognition. rLCMV-GP1-FLAG was generated by insertion PCR utilising Q5 site-directed mutagenesis kit (New England BioLabs) and FLAG primers, according to the manufacturer’s instructions. Positive colonies were verified via colony PCR and DNA sequencing (Genewiz).

### Recovery of rLCMV-WT, rLCMV-eGFP and rLCMV-GP1-FLAG

Recovery of infectious variants of LCMV has previously been described [32], based on the work of the de la Torre group [62]. In brief, six-well plates were seeded with 2 x 10^5^ BSR-T7/well 1 day prior to transfection in 2 mL of Dulbecco modified Eagle medium (DMEM) supplemented with 2.5% fetal bovine serum (FBS), 100 U/mL penicillin and 100 μg/mL streptomycin (2.5% DMEM). After 24 h, the cells were transfected with 1.6 μg pUC57-S-WT, 1.6 μg pUC57-NP (+), 2.8 μg pUC57-L, 2.0 μg pUC57-L (+) and 0.6 μg pCAG-T7pol, combined with 2.5 µL of Mirus TransIT-LT1 transfection reagent (Mirus Bio) per μg of DNA, in 200 µL of Opti-MEM. For rLCMV-eGFP recovery, the WT plasmid was replaced with pUC57-S-eGFP. A control sample, in which pUC57-L and pUC57-L (+) was omitted, was included for each virus recovery experiment. At 24 hours post transfection (hpt), media containing the transfection mix was removed and replaced with fresh 2.5% DMEM. Reinfection of fresh monolayers was carried out in six-well plates seeded with 2 × 10^5^ BHK cells/well 1 day prior to infection in 2 mL of DMEM supplemented with 2.5% FBS, 100 U/mL penicillin and 100 μg/mL streptomycin. Fresh BHK cells were washed with phosphate-buffered saline (PBS) twice prior to infection. Supernatants from transfected BSR-T7 cells were collected at 120 hpt, centrifuged at 4000 *x g* for 15 min and 1 mL used to infect fresh BHK cells in a 6-well plate in DMEM with 2.5% FBS. Cell supernatant from infected BHK cells was collected at 72 hours post infection (hpi), centrifuged at 4000 *x g* for 15 min and viral stocks were stored as aliquots at -80°C. This protocol was followed for successful recovery of the newly developed mutant rLCMV-GP1-FLAG, with the S plasmid of LCMV being replaced with the recently generated pUC57-RQ-S-FLAG. The viral supernatant gained from successful infection following transfection was centrifuged at 4000 *x g*, aliquoted (80 µL) and frozen for subsequent viral titration and bulking.

### Virus infections

For the generation of a bulk stock of rLCMV-GP1-FLAG, a T175 flask was seeded with 5 x 10^6^ BHK-21 cells one day prior to infection at an MOI 0.001. At 3 days post infection, viral supernatant was harvested and centrifuged at 4000 *x g* to remove cell debris, aliquoted (80 µL) and frozen for subsequent viral titration. For viral infections, cell monolayers were infected with LCMV at the specified MOI in either serum-free (SFM), 2.5% or 10% FBS DMEM, depending on cellular requirements, at 37°C. After 1 h, the inoculum was removed and SFM, fresh 2.5% or 10% DMEM was then applied for the duration of the infection. For synchronised infections, LCMV incubated on ice for 1 h to facilitate adsorption. Subsequently, the inoculum was removed, monolayers washed three times with PBS and SFM, fresh 2.5% or 10% DMEM was then applied for the duration of the infection.

### Viral titration

Virus titers were determined by focus forming assays. Viral stocks requiring titration were serially diluted in SFM to infect fresh monolayers of BHK cells seeded at 1 x 10^5^ in a 24 well-plate and incubated at 37°C for 1 h. After infection, medium containing virus was removed and 2 mL of overlay containing 10% FBS DMEM and 1.6% methylcellulose at a 1:1 ratio was reapplied to cells, then incubated for a further 3 days at 37°C. For rLCMV-eGFP titration, whole-well images were taken using an Incucyte S3 live cell imaging system (Sartorius) and rLCMV-eGFP foci were then counted and virus titers were determined as focus forming units/mL (FFU/mL). For rLCMV-WT titration, cells were fixed using 4% (vol/vol) paraformaldehyde (PFA) for 15 min and washed three times with PBS. Cells were then incubated with permeabilisation buffer (0.1% [vol/vol] Triton X-100, 2% [wt/vol] FBS in 1 x PBS) for a further 15 min and washed three times with PBS. After permeabilisation, cells were incubated with 1 mL blocking buffer (2% [wt/vol] FBS in PBS) for 1 h, then incubated for 1 h with 150 µL/well LCMV NP primary antibody (in-house, 1:1000 in blocking buffer), and washed three times with PBS. Following this, cells were incubated for 1 h with 594 Alexa Fluor secondary antibody (Life Technologies; 1:500 in blocking buffer) and washed four times with PBS. The Incucyte S3 live cell imaging system (Sartorius) was then used to image whole wells of the plate to detect red rLCMV-WT foci, which were counted, and virus titers determined. For rLCMV-FLAG titration, cells were also stained with 150 µL/well FLAG (Sigma, 1:500 in blocking buffer) primary antibody and 488 (FLAG) Alexa Fluor secondary antibody (Life Technologies; 1:500 in blocking buffer), then washed and imaged as described above.

### Reverse transfection of siRNA library

Trypsinised SH-SY5Y cells in 10% FBS DMEM were counted using a hemocytometer and used to make a cell suspension containing 1 x 10^5^ cells/mL. A master mix was made resulting in 0.3 µL of Lipofectamine RNAiMAX reagent (Invitrogen) and 16.7 µL of Opti-MEM per well and 17 µL of this master mix was pipetted into each well of a 96-well plate. A 3 µL volume of working stock siRNA (1 μM) was pipetted into the transfection master mix and mixed, resulting in a final concentration of 3 pmol of siRNA per well. Transfection master mix and siRNA were incubated for 20 min, following which 1 × 10^5^ cells in 10% DMEM (100 µL total volume) was then applied per well. Cells were incubated with the transfection and siRNA mix for 24 h at 37°C, following which 60 µL of the medium was removed and replaced with 200 µL of fresh 10% FBS DMEM to dilute out any potential toxic effects of the siRNAs or transfection reagent. At 6 h post dilution, the medium was removed and cells were washed in PBS prior to infection with rLCMV-eGFP at an MOI of 0.2 in 100 µL of 10% FBS DMEM for 1 h. Following infection, virus was removed and cells were washed twice with PBS to remove any unbound virus, then supplemented with 200 µL 10% FBS DMEM. At 24 hpi, eGFP fluorescence intensity was determined using the Incucyte S3 live cell imaging software as a measure of virus multiplication. Following initial imaging, 200 µL viral supernatant for each siRNA was transferred onto fresh SH-SY5Y cells seeded previously at 1 x 10^4^ cells/mL. At 24 hpi, the eGFP fluorescence intensity was determined using the Incucyte S3 live cell imaging software as a measure of infectious virion production. The total integrated intensity of eGFP (TIIE; green count units [GCU] × μm^2^/image) was first normalised to confluency per well and then analysed as a percentage of the total green integrated intensity in positive control wells containing virus and lipofectamine, but omitting siRNA. Normalised values were averaged between four experimental repeats.

### Immunofluorescence (confocal microscopy)

Trypsinised A549 cells were seeded onto a 19-mm round glass coverslip (VWR) in a 12-well plate at 1 x 10^5^ cells/well, followed by incubation at 37°C. After 14 h, BFA was added to cells (5000 ng/mL) for 1 h prior to fixation. For IF (omitting AP4E1, see below), cells were washed twice in PBS prior to fixation in 4% (vol/vol) paraformaldehyde in PBS for 15 min at room temperature. After fixation, the cells were washed three times in PBS and then incubated in permeabilisation buffer (0.1% [vol/vol] Triton X-100, 1% [wt/vol] bovine serum albumin [BSA] in PBS) for 15 min at room temperature. Following permeabilisation, the monolayers were washed three times with PBS and incubated with blocking buffer (1% [wt/vol] BSA in PBS) for 1 h. Subsequently, primary antibody was diluted in BSA blocking buffer as follows: NP (1:500, in house [sheep]), FLAG (Sigma; 1:250 [mouse]) and COPA primary antibody (GeneTex; 1:100 [rabbit]) and incubated for 1 h at room temperature. The cells were then washed three times with PBS and incubated with corresponding Alexa Fluor 488, 594 and 647 secondary antibodies (Life Technologies; 1:500 in BSA blocking buffer) for 1 h at room temperature in a light protected vessel. Cell monolayers were then washed three times with PBS and mounted onto glass coverslips with the addition of Prolong Gold Antifade reagent with DAPI (Thermo Fisher Scientific), cured, sealed and stored at 4°C. Images were then taken on an LSM 880 confocal microscope (Zeiss) and processed using Zen (Blue Edition) software and Fiji (Image J). Line scan analysis was performed utilising Zen (Blue Edition).

For AP4E1 IF, cells were washed twice with PBS and fixed with 100% methanol for 5 min on ice. After fixation, the cells were washed three times in PBS and then incubated with blocking buffer (0.1% [wt/vol] Saponin, 1% [wt/vol] BSA in PBS) for 30 min. Subsequently, primary antibodies diluted in saponin/BSA blocking buffer containing NP (1:500, in house [sheep]), FLAG (Sigma; 1:250 [rabbit]) and AP4E1 primary antibody (BD Biosciences; 1:75 [mouse]) was incubated for 1 h at room temperature. The cells were then washed three times with PBS and incubated with corresponding Alexa Fluor 488, 594 and 647 secondary antibodies (Life Technologies; 1:500 in saponin/BSA blocking buffer) for 1 h at room temperature in a light protected vessel. Cell monolayers were then washed three times with PBS and mounted onto glass coverslips with the addition of Prolong Gold Antifade reagent with DAPI (Thermo Fisher Scientific), cured, sealed and stored at 4°C. Images were then taken on an LSM 880 confocal microscope and processed as described above.

### Inhibition of retrograde transport

Trypsinised A549 cells were seeded into 24-well plates at 1 x 10^5^ cells/well and incubated at 37°C. After 16 to 24 h, the cells were pre-treated with BFA at the indicated concentration for 45 min in SFM. Following this, rLCMV, rLCMV-eGFP, A/WSN/33 H1N1 (denoted as rIAV herein) and rIAV-eGFP was added directly to each well at an MOI of 0.1. For WT viruses, 24 hpi cells were lysed for analysis via western blotting. Densitometry was performed using Fiji software and NP expression normalised to confluency per well, then analysed as a percentage of virus only DMSO control. Normalised values were averaged between three biological repeats for WT viruses (n=3). For eGFP viruses, at 24 hpi the Incucyte S3 live cell imaging system (Sartorius) was utilised to measure eGFP fluorescence. The total integrated intensity of eGFP (TIIE; green count units [GCU] × μm^2^/image) was first normalized to confluency per well and then analysed as a percentage of the total green integrated intensity in virus only DMSO control. Normalized values were averaged between three biological repeats for eGFP mutant virus (n=3). For titration following BFA treatment, viral supernatant was collected for mock, virus, 50, 500 and 5000 ng/mL. Focus forming assays were performed for each BFA condition as previously described for LCMV-eGFP (n=3).

### Quantitative PCR

A549 cells were seeded into a 12-well plate at 2 x 10^5^ cells/well. Following 16-24 h cells were infected at an MOI of 1 with rLCMV-WT in minimum volume 10% FBS DMEM media (400 µL) for 1 h. Subsequently, monolayers were supplemented with an additional 600 µL 10% FBS DMEM and infection was allowed to progress for a further 24 h. At 24 hpi, cell monolayers were washed twice with PBS and trypsinised with 500 µL trypsin. Following cell detachment, 500 µL 10% FBS DMEM was added to each well. Subsequently, cells were pelleted by centrifugation at 500 *x g* for 1 min. RNA extraction was performed following the manufacturer’s instructions for Monarch Total RNA Miniprep Kit (New England BioLabs). Two step qPCR was carried out firstly by converting RNA into cDNA following the manufacturer’s instruction for LunaScript RT SuperMix Kit (New England BioLabs). Subsequently, qPCR was performed using the cDNA following the manufacturer’s instructions for Luna Universal qPCR Master Mix. Analysis of mRNA expression via qPCR for mock and virus (MOI 1) for the genes of interest; AP4B1, AP4E1, AP4M1, COPA, COPB1, COPB2, LCMV NP and GAPDH were performed. The primer sequences used were as follows: AP4B1 (5’ -CTG GCG TTA CGG AGC ATG T-3’ and 5’ -GAC CAT TGA GAA TAG GCT GTT GT-3’), AP4E1 (5’ -TTC ATG CAA TCA AGT TAG CCC A-3’ and 5’ -TCA GTG CCA TAC ACA CTT CTA CT-3’), AP4M1 (5’ -CTT CCT TCC TAG CGG CTC TG-3’ and 5’ -GAT TCC TGG CCC ATA ACC TCT -3’), and 5’-CAG TGT CAT TAA CTC CAA CCA CA-3’), COPA (5′-CCA CTA TCA GAA TGC CCT ATA CC-3′ and 5′-CCA CAA ACC CAT CTT CAT CC-3′), COPB1 (5′-ACA GAGA GAA AGA GGC AGC AGA-3′ and 5′-GCA AGG TATA CAC TGG TTT GGT TC-3′), COPB2 (5′-GTG GGG ACA AGC CAT ACC TC 3′ and 5′- GTG CTC TCA AGC CGG TAG G-3′), GAPDH (5’ -AGG GTC ATC ATC TCT GCC CCC-3’ and 5’ -TGT GCT CAT GAG TCC CAC GAT-3’) and LCMV NP (5’ -TGT GGC AGA ATG TTG TGA AC-3’ and 5’ -AAA AGA AGA AAG AGA TCA CCC C-3’). Prior to qPCR, primer efficiencies were performed to ensure primer sets were between 80-100% amplification efficiency. The mRNA expression of LCMV NP was calculated for mock and MOI 1 to ensure changes in cellular gene mRNA expression was a result of infection. Ct values were normalised for sample variation in GAPDH, and fold change calculated and appropriate NTC controls used. Each individual gene fold change value for MOI 1 RNA isolates were then normalised to assess up/down regulation in mRNA expression. Relative percentage change values were averaged between four experiment repeats.

## Supporting information

supplemental siRNA dataset

## ACKNOWLEDGEMENTS.

We acknowledge funding from MRC grant MR/T016159/1 in support of JNB, JF, AS and HNT, BBSRC PhD studentship grant to AS, UKHSA PhD studentship grant to OB, and Wellcome trust studentship 102174/B/13/Z to EJAAT. BAR was supported by EU Marie Skłodowska-Curie Actions (MSCA) Innovative Training Network (ITN) H2020-MSCA-ITN-2016, grant agreement 721367 (HONOURS). The authors thank Dr Ruth Hughes and Dr Sally Boxall of the bioimaging facility, Faculty of Biological Sciences, University of Leeds, for their expert assistance and use of the Zeiss LSM880 confocal microscope, funded by Wellcome Trust grant WT104918MA. We also acknowledge The Wellcome trust equipment grant 221538/Z/20/Z, which supports the use of the Incucyte live cell imaging platform.

## SUPPLEMENTARY FIGURE LEGENDS

**Figure S1.**
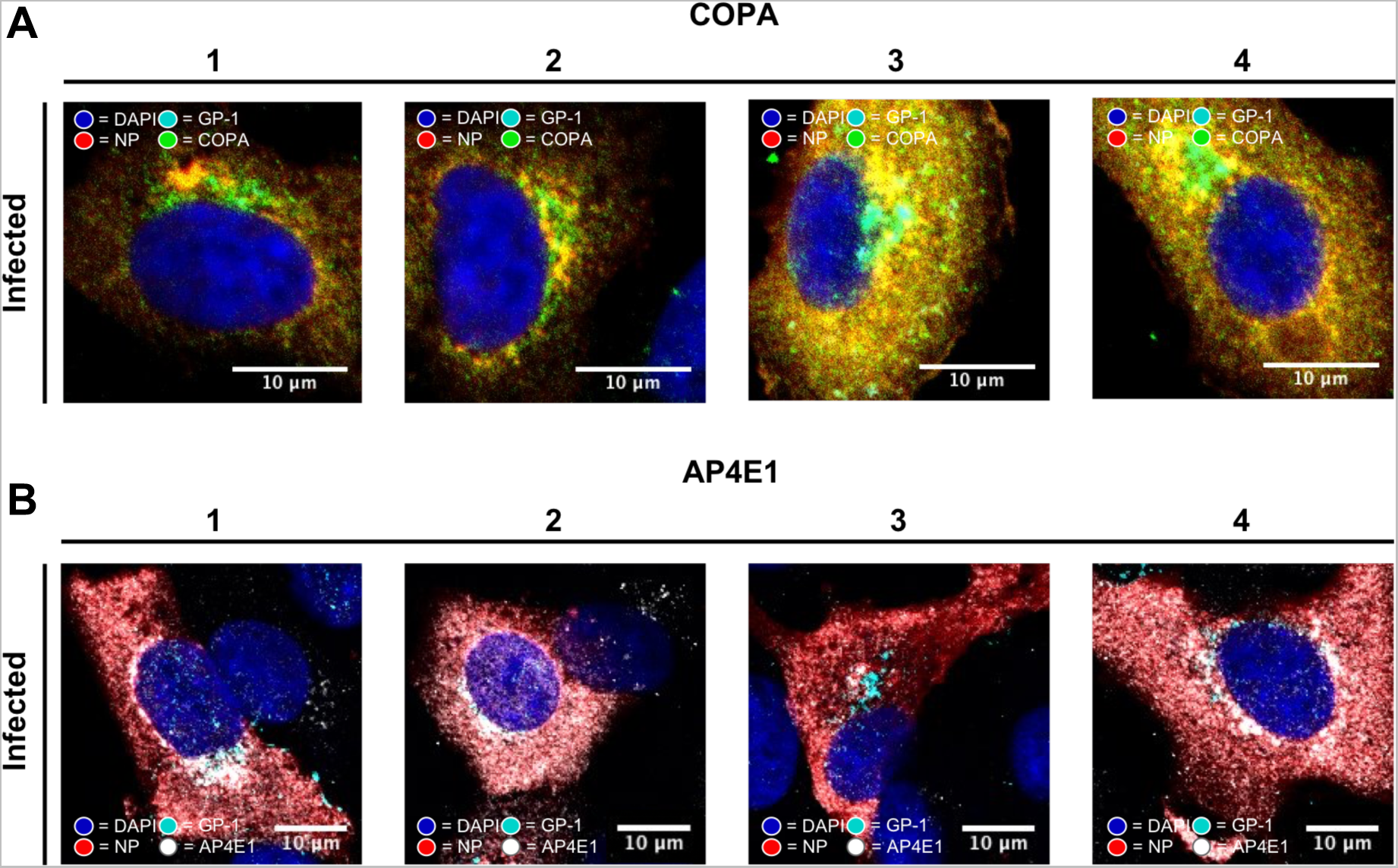
Co-localization of LCMV components with COPA and AP4E1. (A) Confocal microscopy of four different A549 cells infected with rLCMV-eGFP and stained at 24 hpi with COPA (green), GP-1 (cyan), and NP (red) and DAPI. (B) Confocal microscopy of four different A549 cells infected with rLCMV-eGFP and stained at 24 hpi stained with AP4E1 (white), GP-1 (cyan), and NP (red) and DAPI.

**Figure S2.**
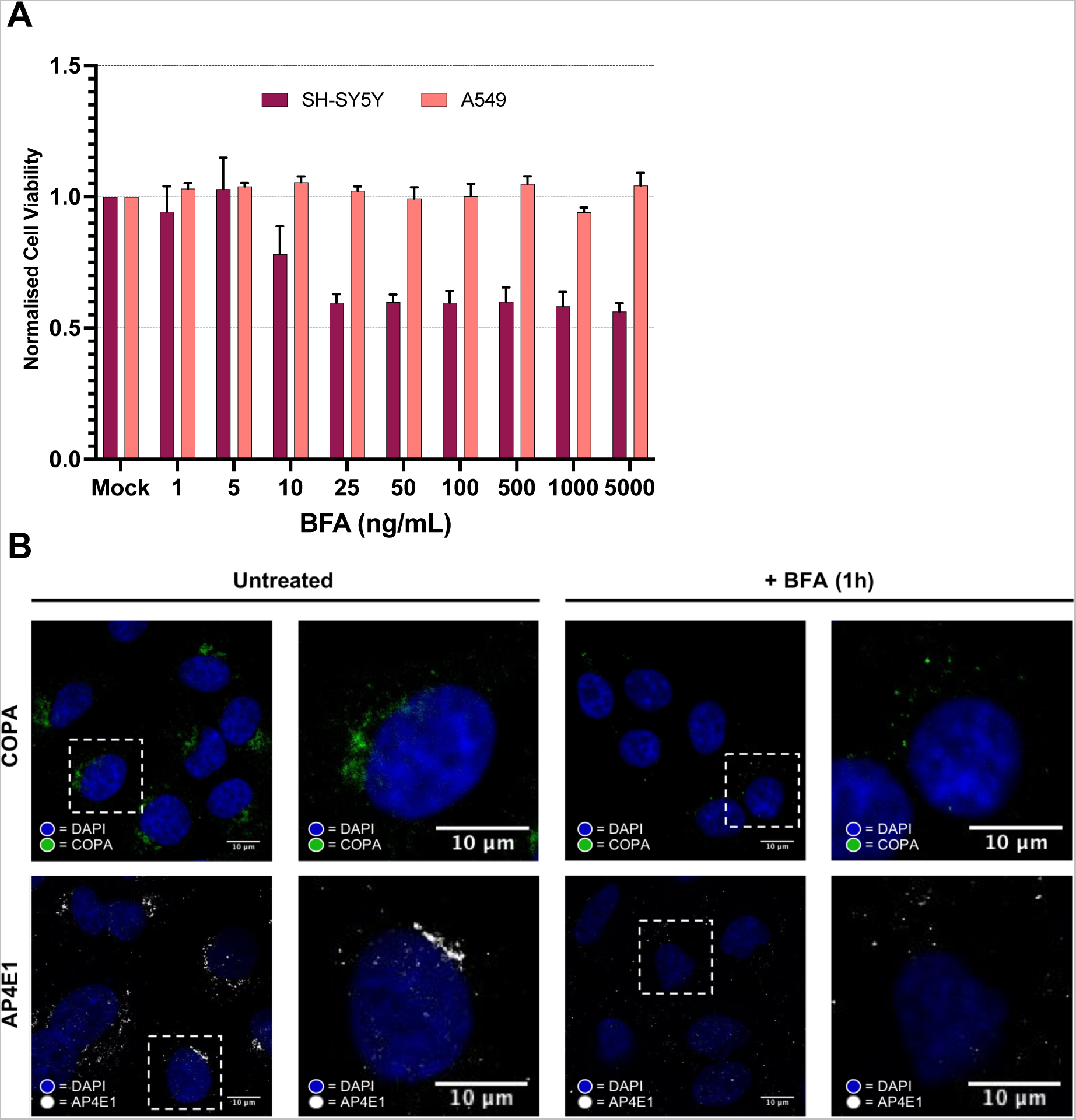
Addition of BFA at late stages of rLCMV-GP1-FLAG infection results in changes in the cellular distribution of COPA and AP4E1. (A) Analysis of BFA toxicity in A549 cells at various concentrations as measured by MTT assay, normalized to mock untreated cells. (B) Confocal microscopy images of untreated and BFA-treated A549 cells (5000 ng/mL) infected with rLCMV-GP1-FLAG at an MOI of 0.1, at 24 hpi. The infected cells were stained with DAPI and using antisera specific for either COPA (green; top panels) or AP4E1 (white, bottom panels). A zoomed image for both untreated and BFA-treated cells is shown corresponding to the white boxed area.

## Notes

### Competing Interest Statement

The authors have declared no competing interest.

